## Supplementary figures and images for "Water-soluble 4-(dimethylaminomethyl)heliomycin exerts greater antitumor effects than parental heliomycin by targeting the tNOX-SIRT1 axis and apoptosis in oral cancer cells"

### Figure S1

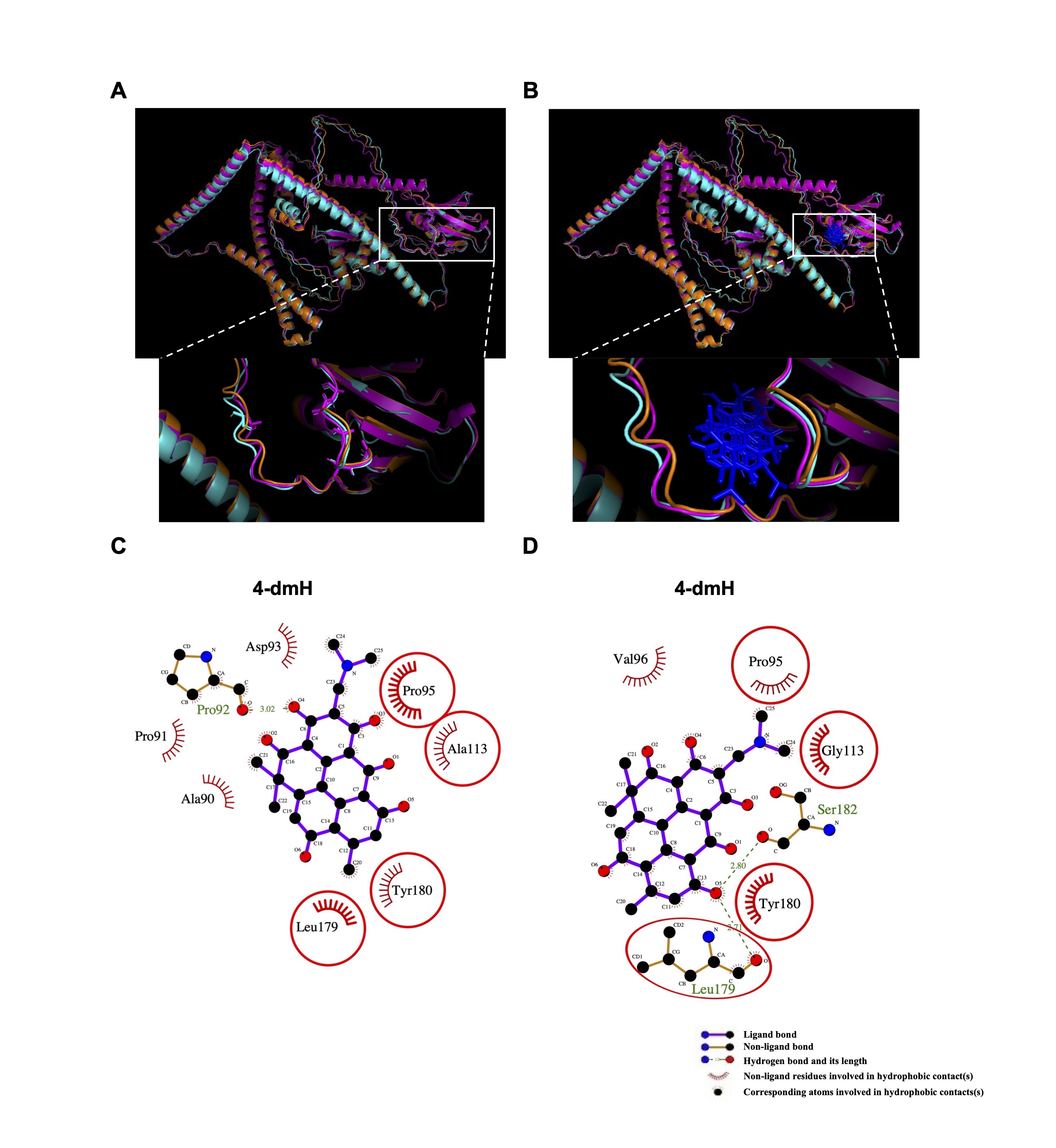
