## Supplementary material for "Water-soluble 4-(dimethylaminomethyl)heliomycin exerts greater antitumor effects than parental heliomycin by targeting the tNOX-SIRT1 axis and apoptosis in oral cancer cells": Figure legends for S1

Figure S1

The simulated tNOX structures (a, b) and the binding modes of 4-dmH after docking study (c, d). (a) Superimposition of three types of tNOX structures, including the original tNOX structure (orange) and the critical residues in tNOX protein substituted with alanine (magenta) or glycine (cyan). The substituted residues were shown as sticks. (b) Superimposition of the docked 4-dmH (blue). (c) Schematic presentations of possible interactions between 4-dmH and the interacted residues in tNOX protein substituted with alanine. (d) Schematic presentations of possible interactions between 4-dmH and the interacted residues in tNOX protein substituted with glycine. The key residues were identified based on the best docking pose of 4-dmH. The red circles and ellipses indicate the identical residues that interacted with different types of tNOX structures.
