## Supplementary material for "Water-soluble 4-(dimethylaminomethyl)heliomycin exerts greater antitumor effects than parental heliomycin by targeting the tNOX-SIRT1 axis and apoptosis in oral cancer cells": Uncrop gel images

Figure 2 (a): SAS

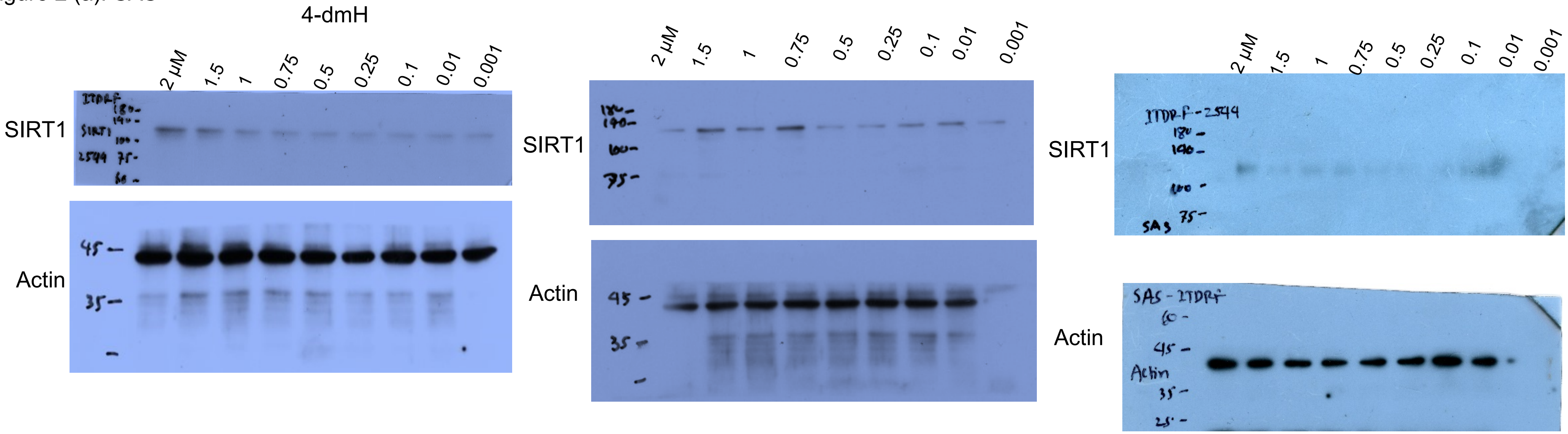


HSC-3

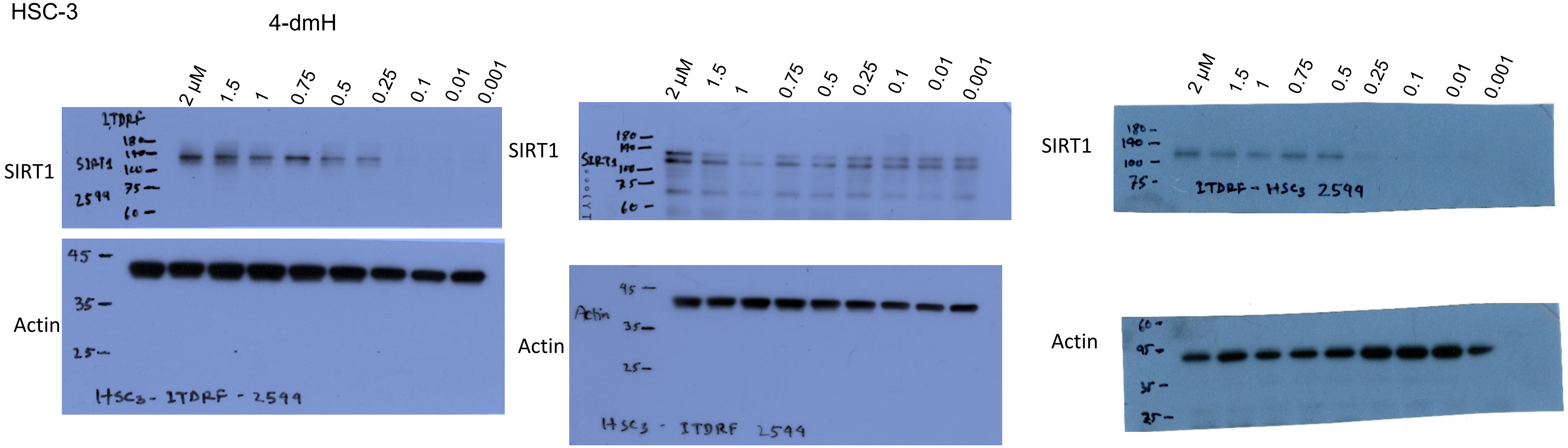

Figure 2 (b): SAS

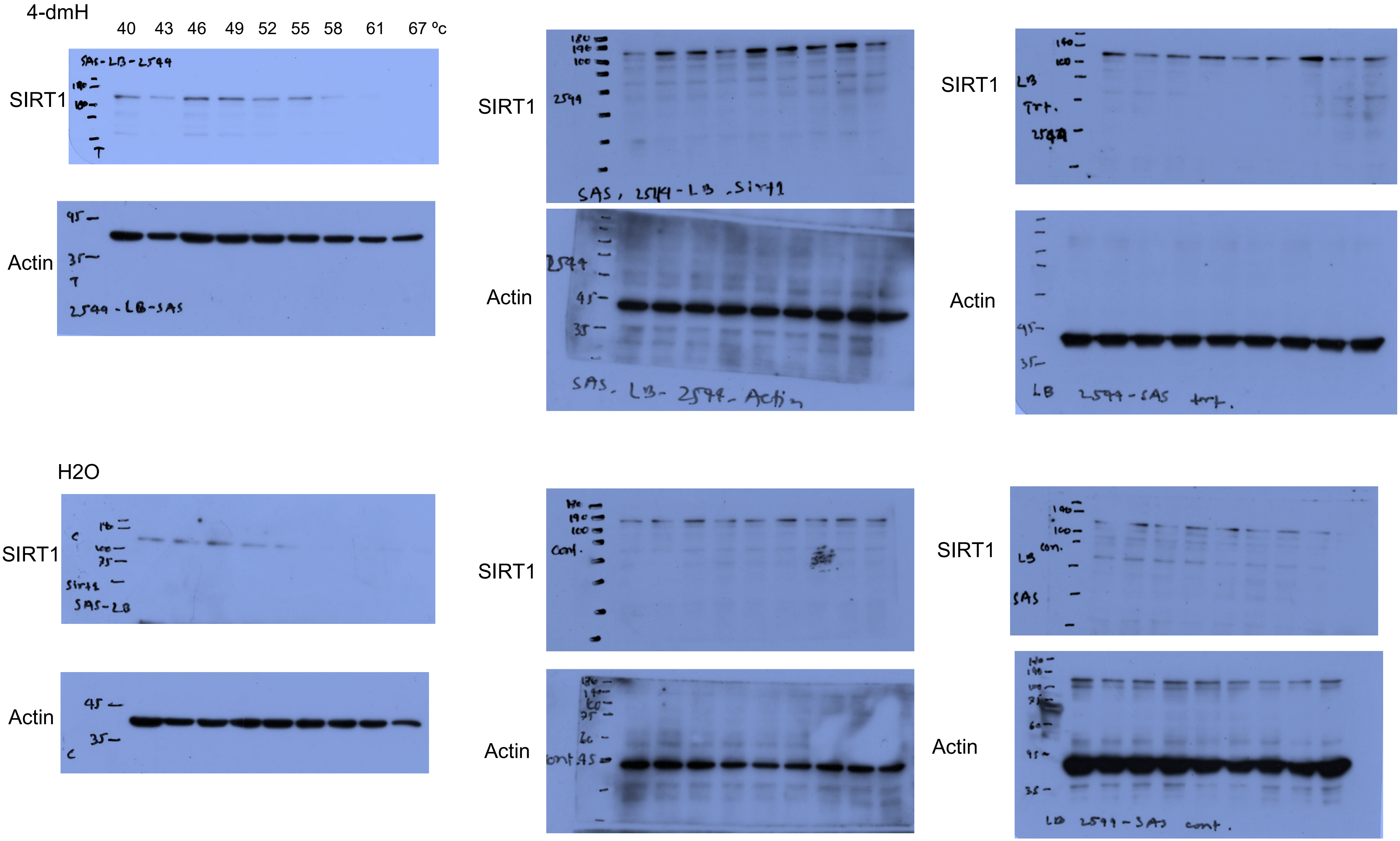

Figure 2 (b): HSC-3

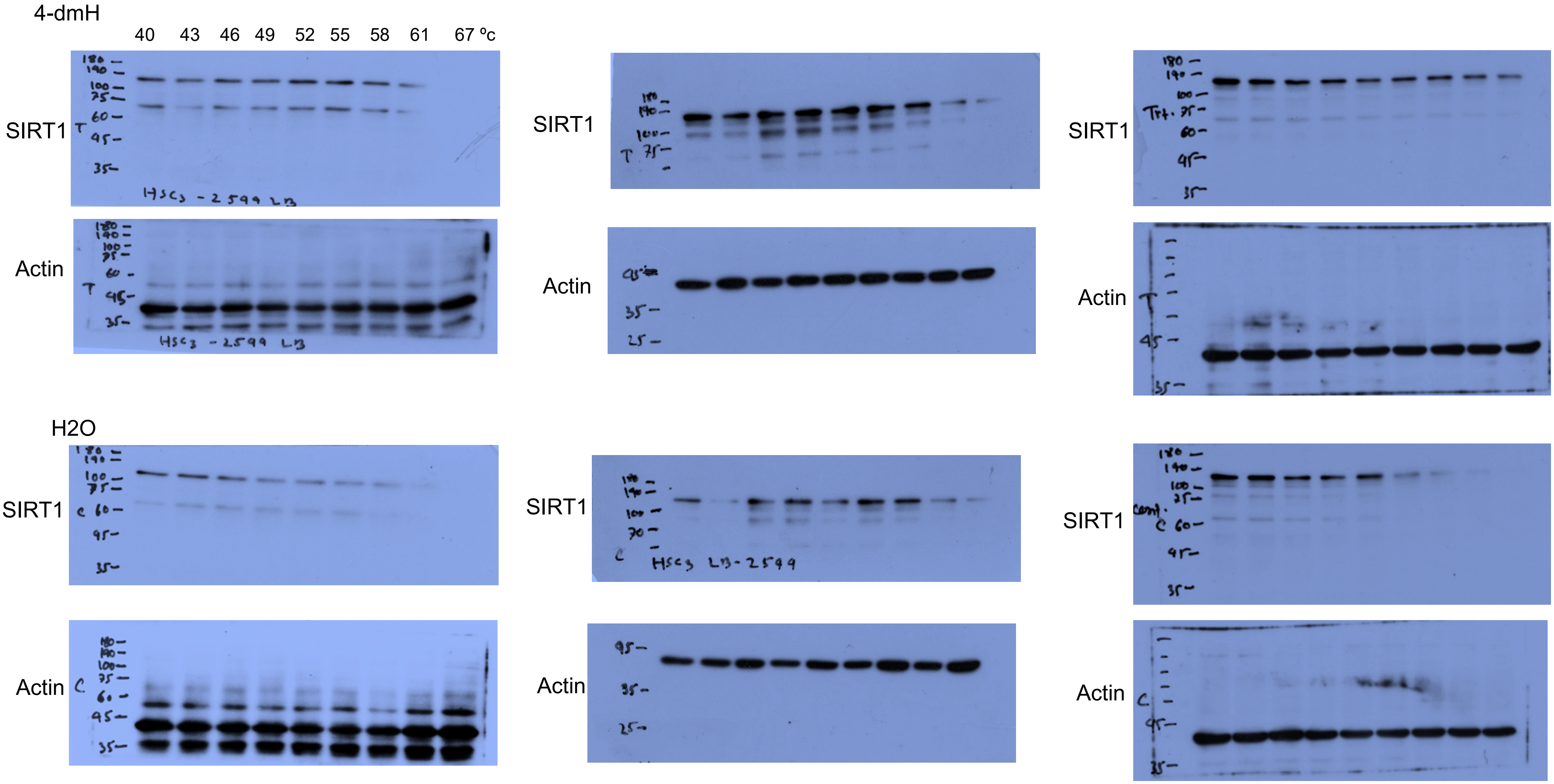

Figure 3 (a): SAS

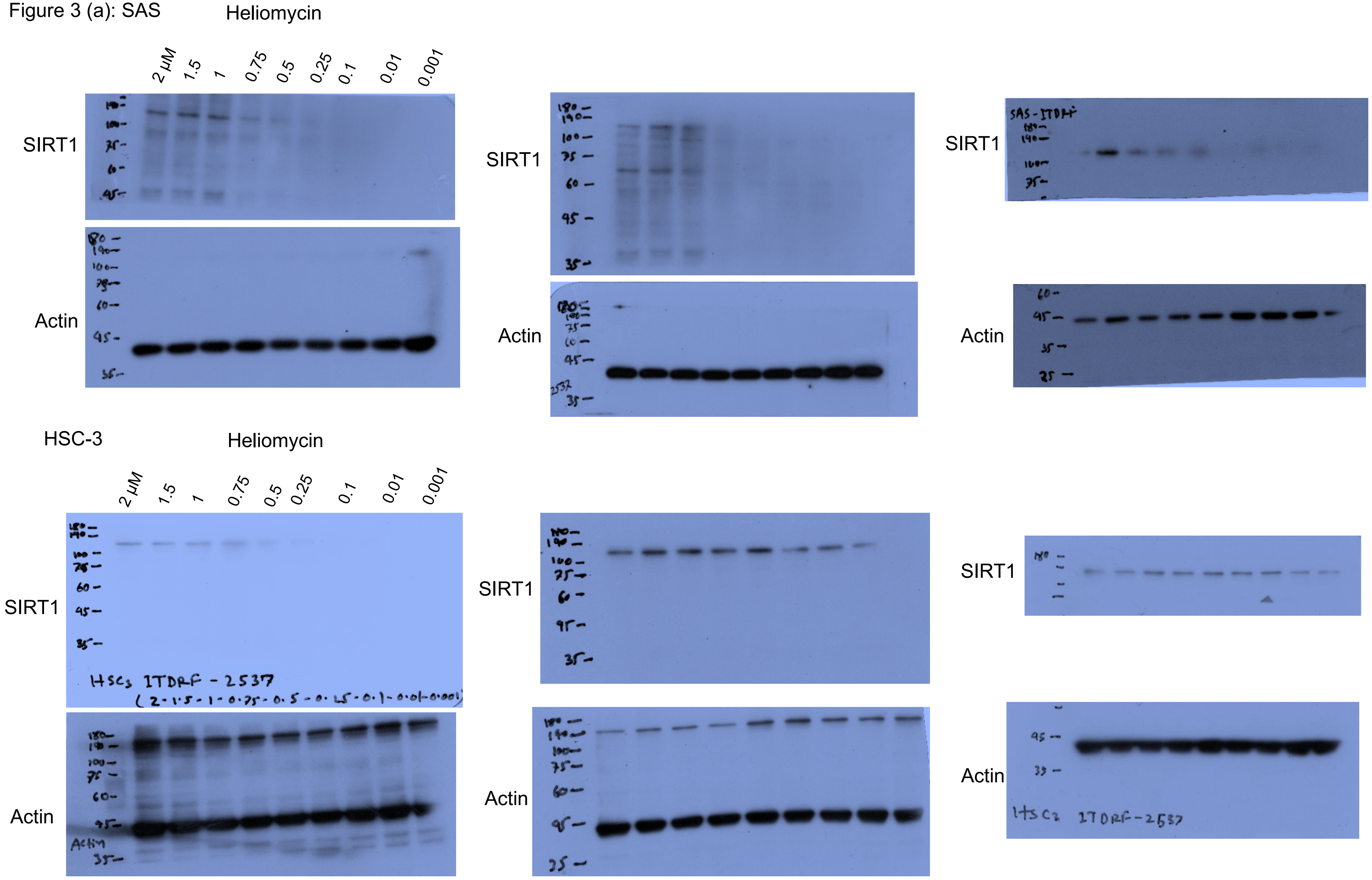

Figure 3 (b): SAS

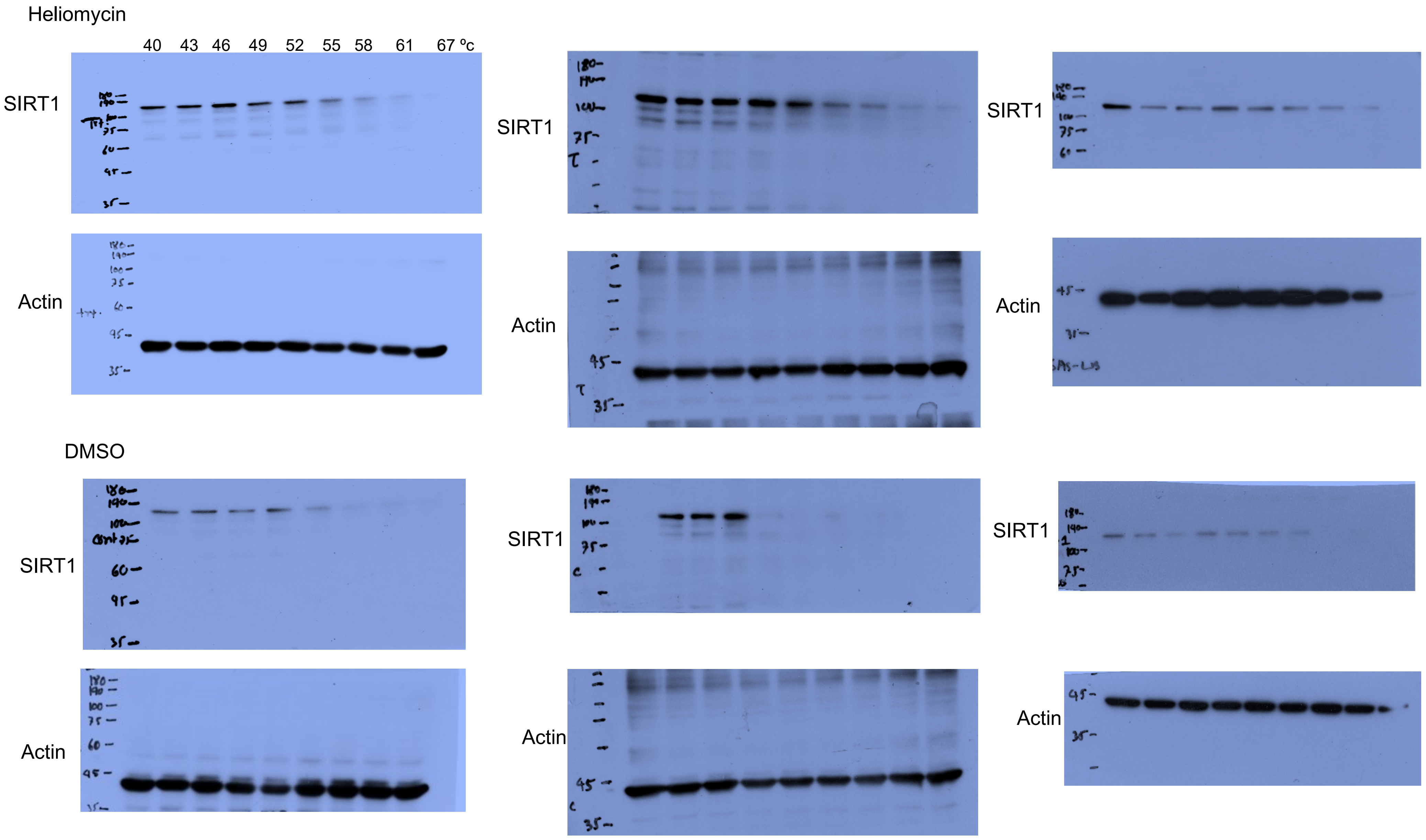

Figure 3 (b): HSC-3

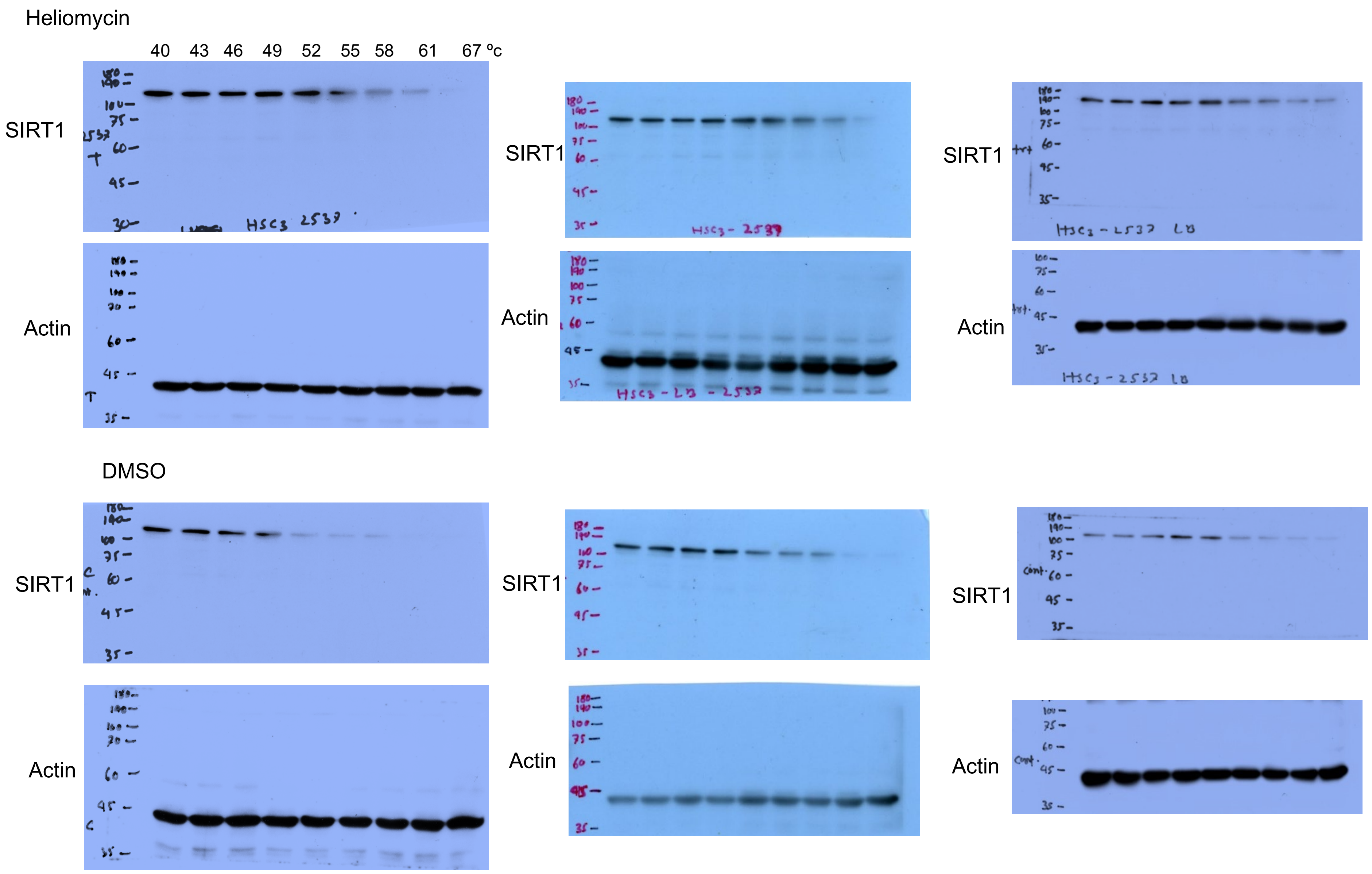

Figure 4 (b): SAS

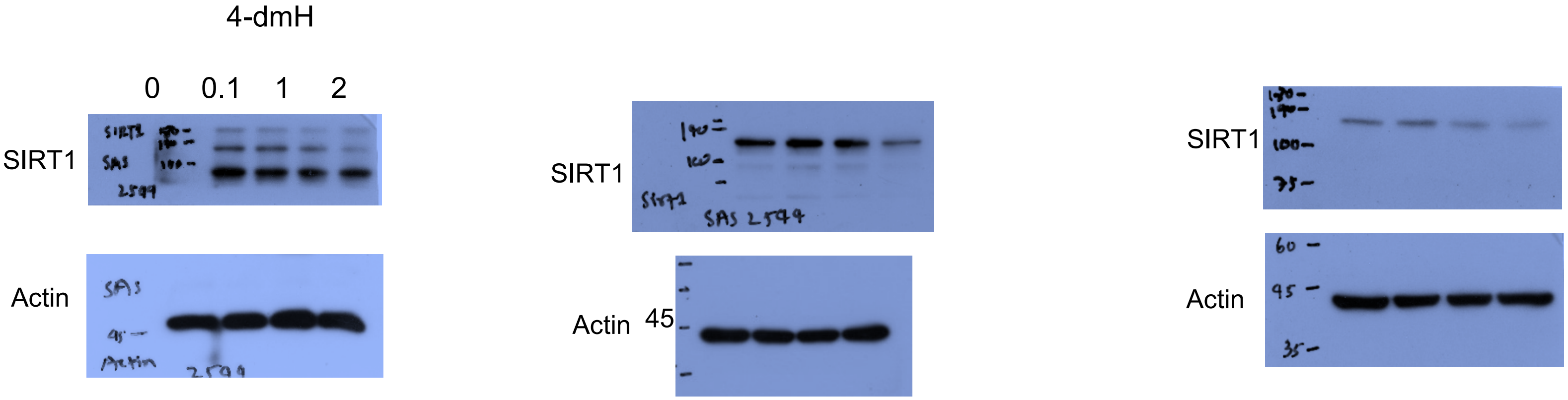


HSC-3

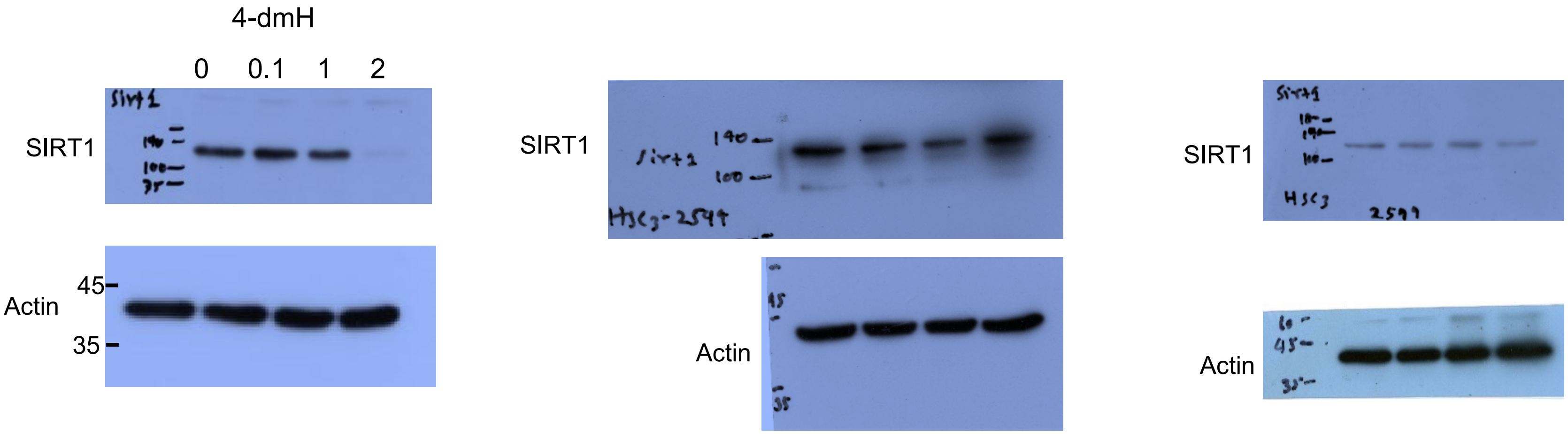

Figure 4 (c): SAS

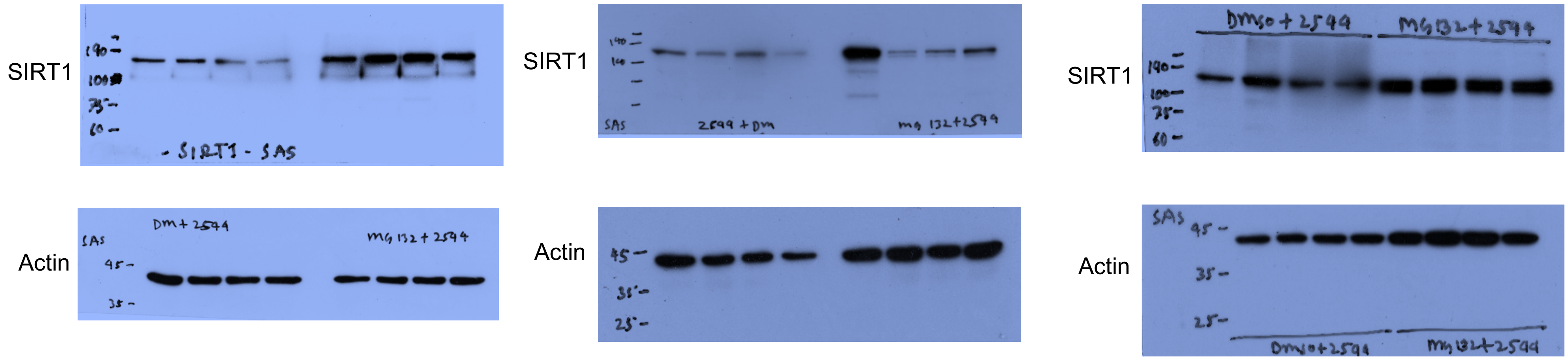


HSC-3

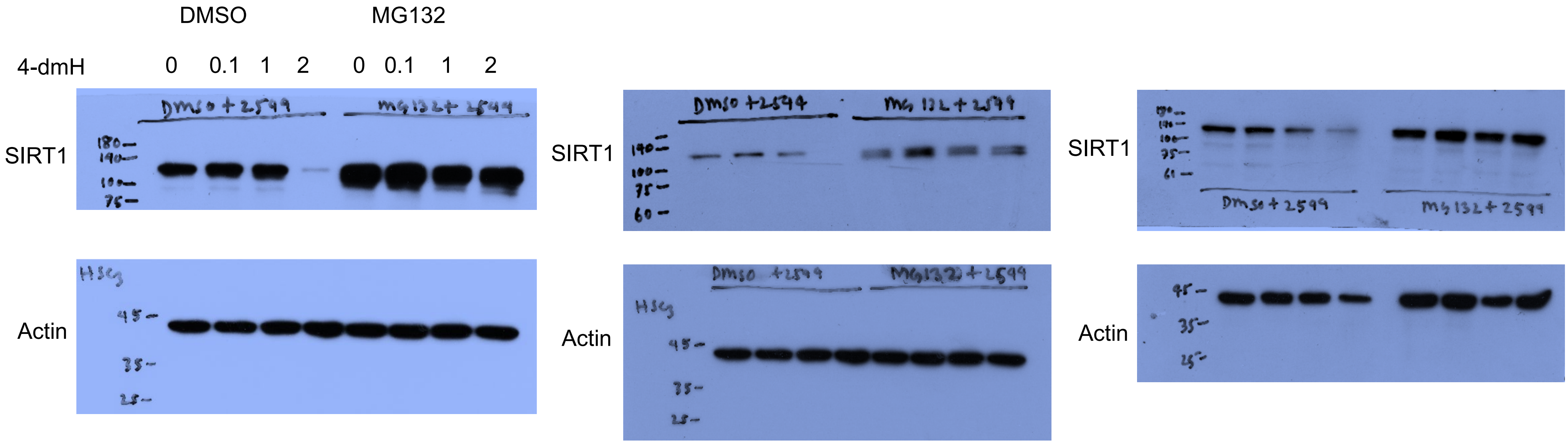

Figure 4 (d): SAS

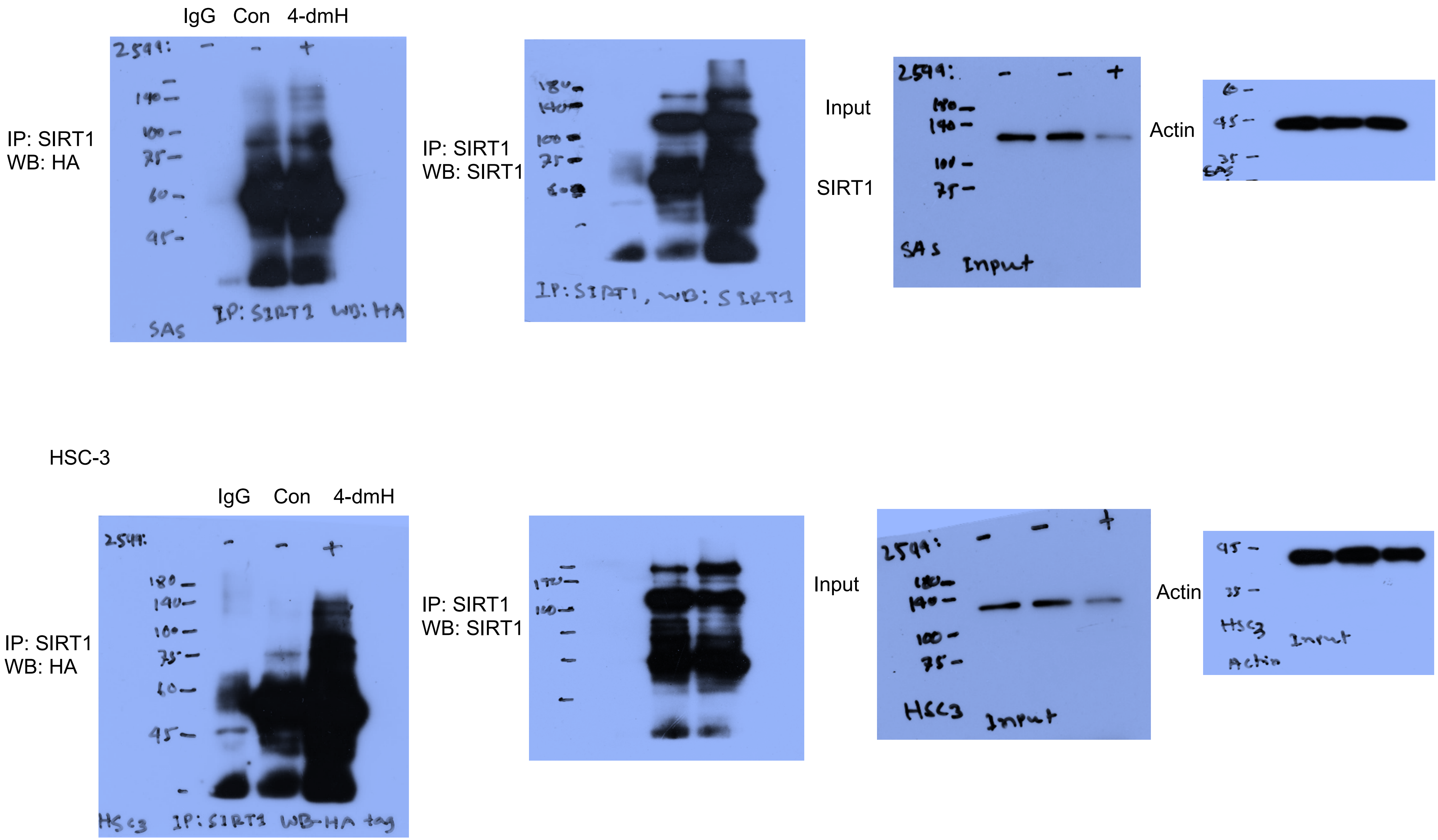

Figure 5 (c): HSC-3

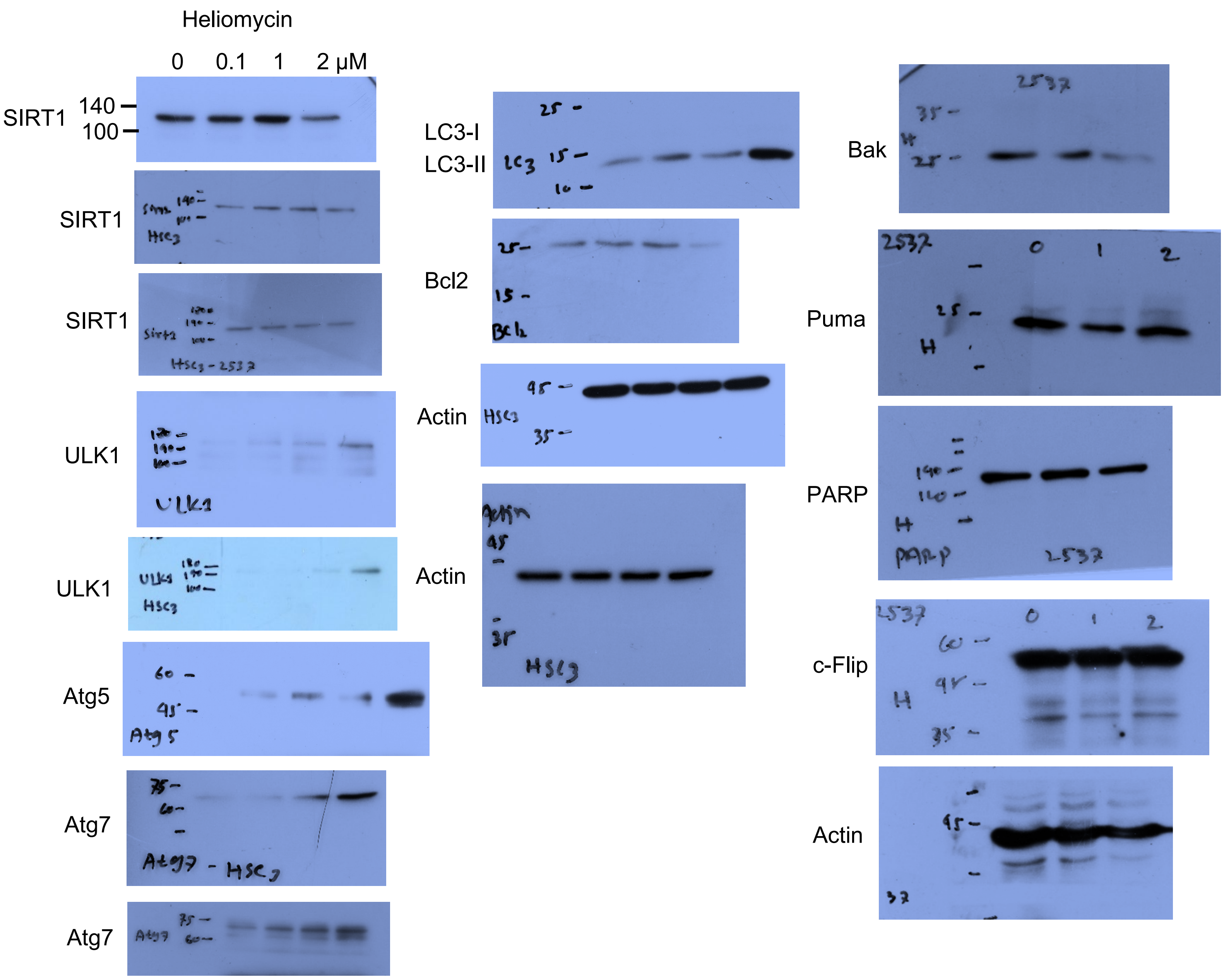

Figure 5 (c): SAS

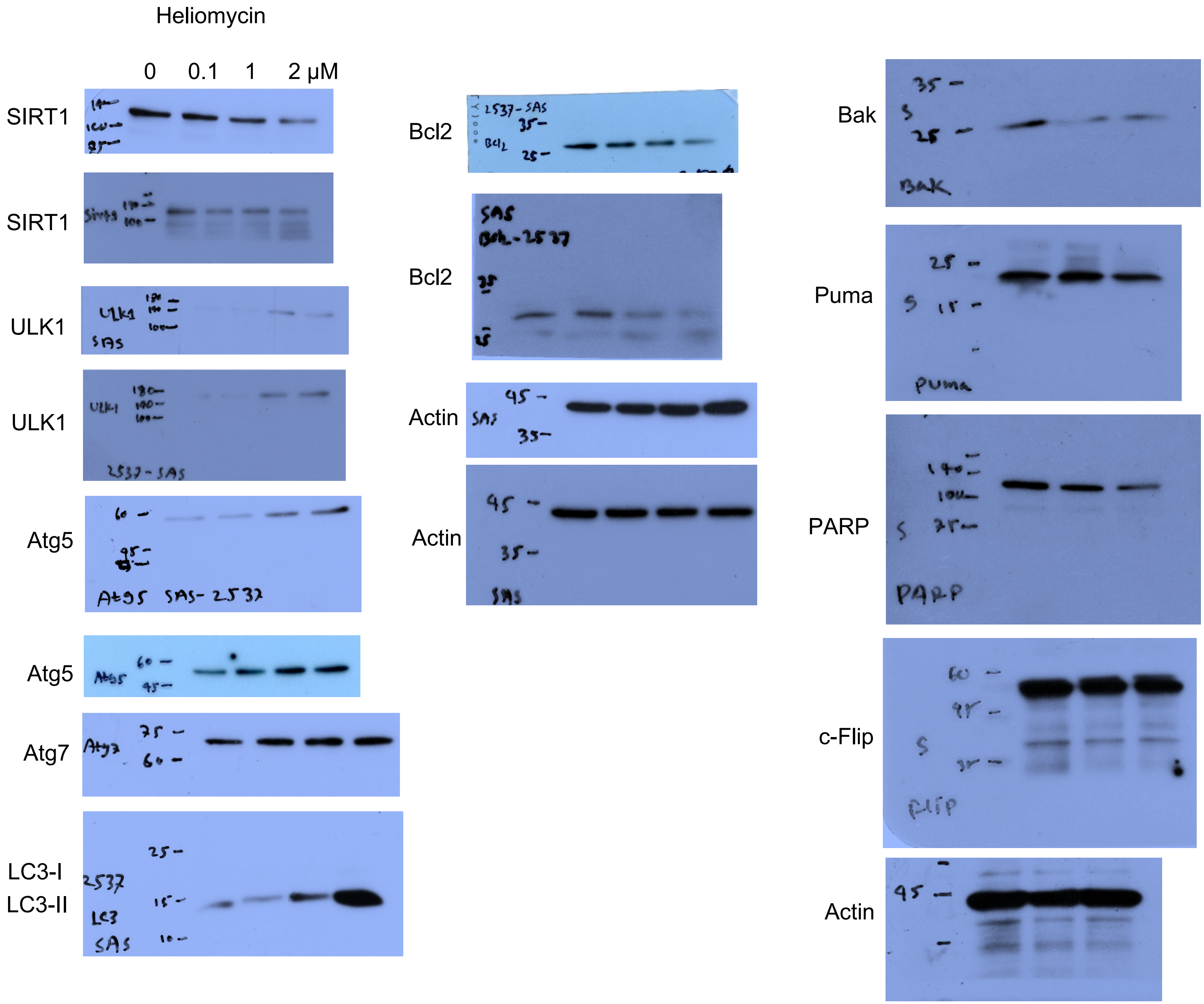

Figure 5 (d): HSC-3

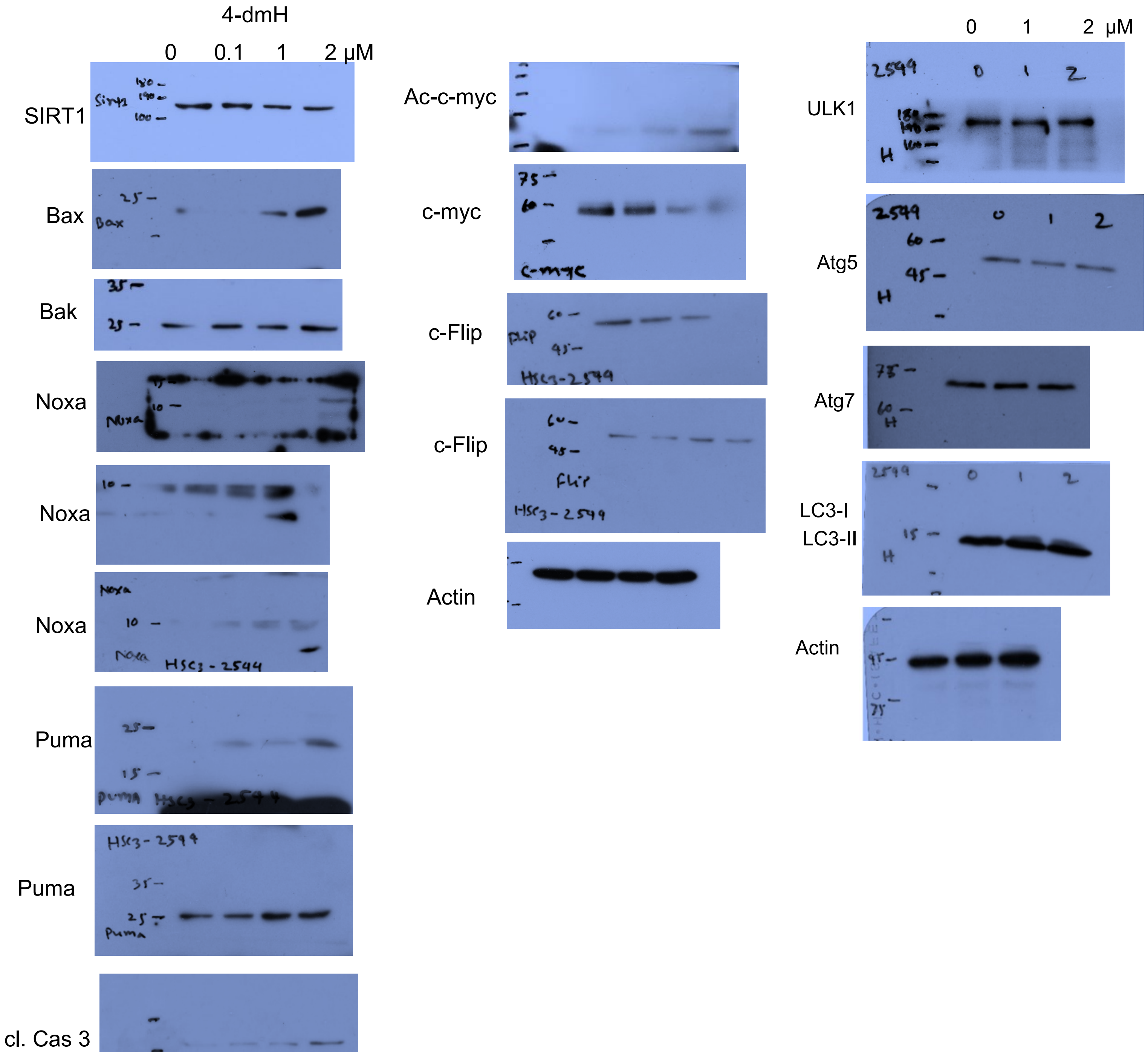

Figure 5 (d): SAS

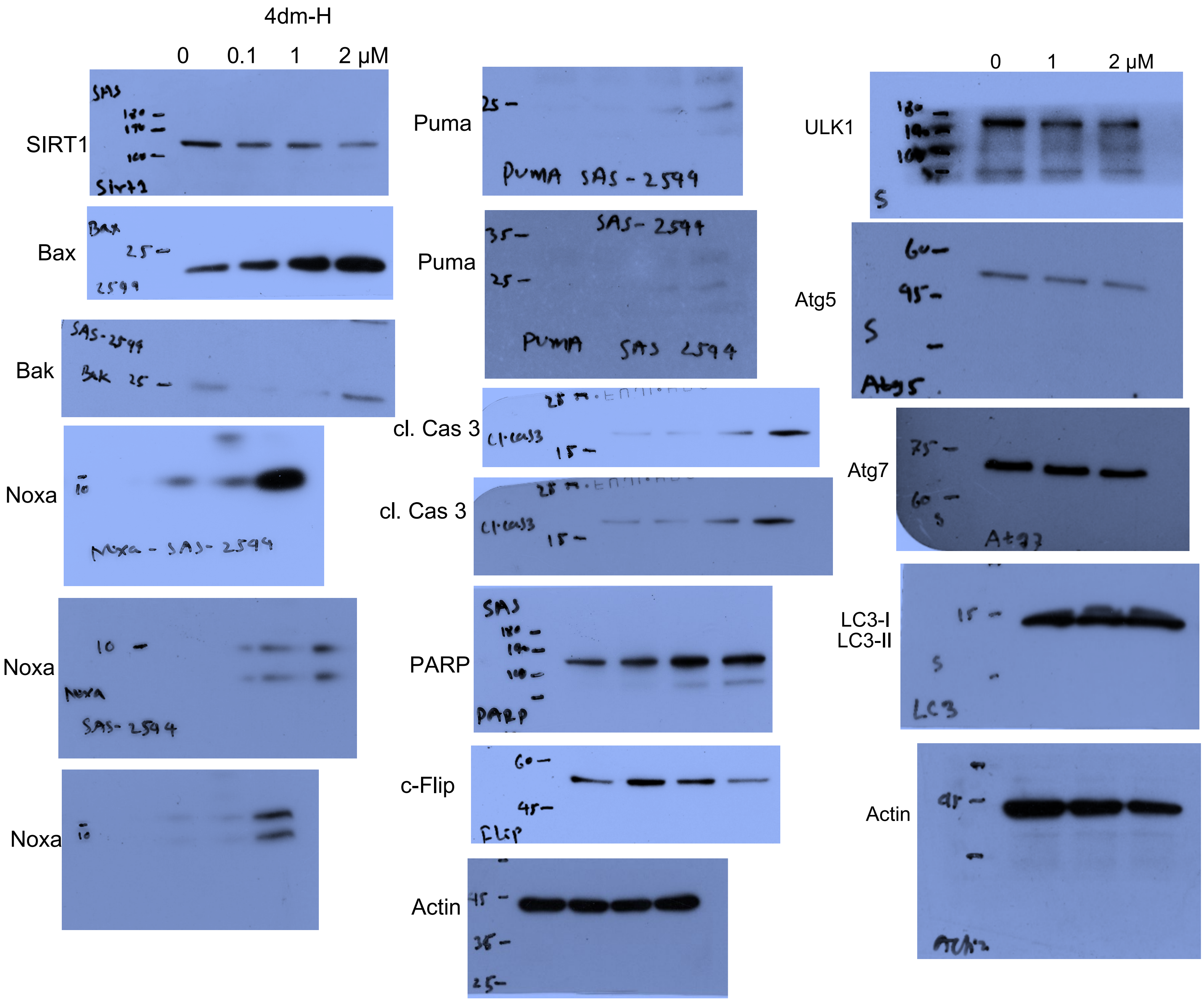

Figure 7 (b): SAS

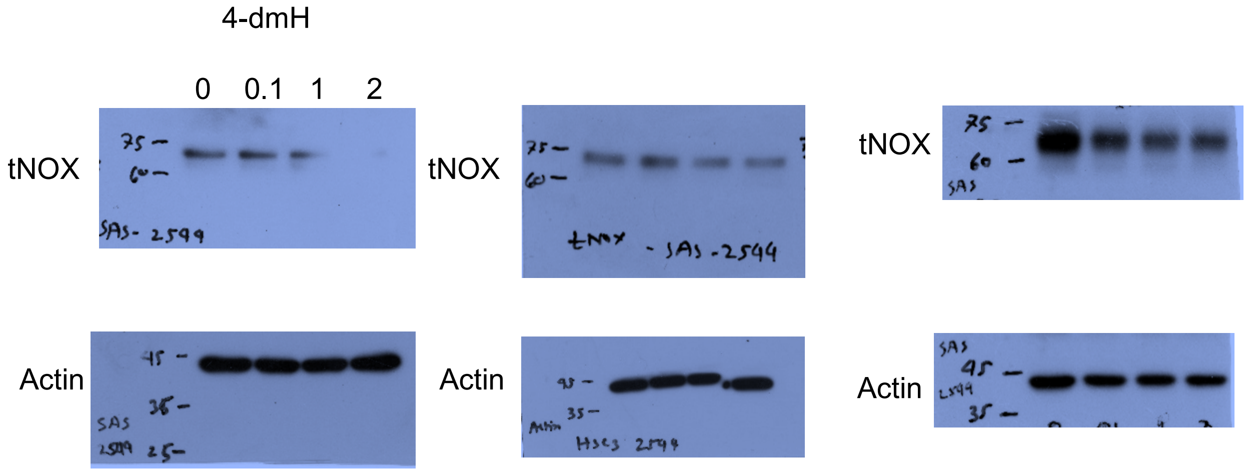


HSC-3

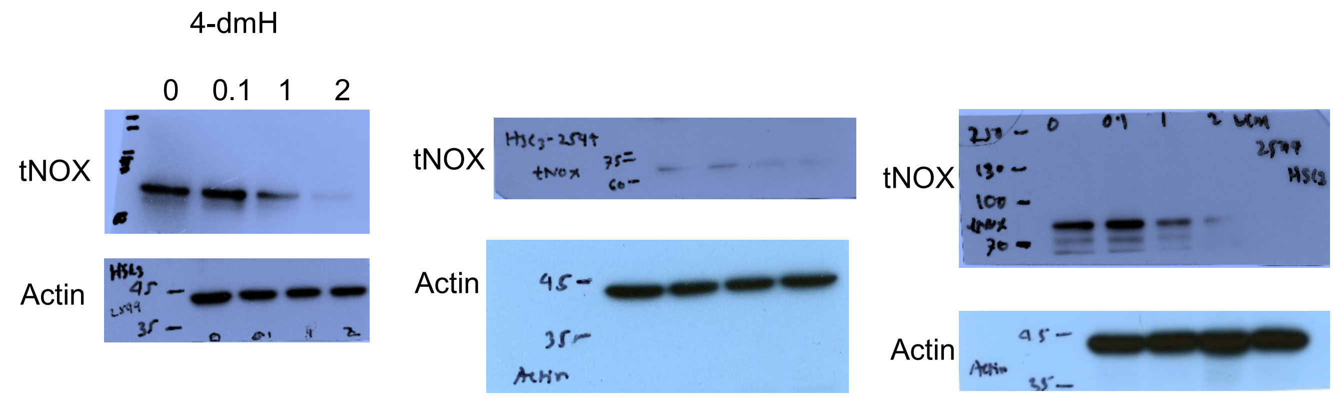


Figure 7 (b): SAS

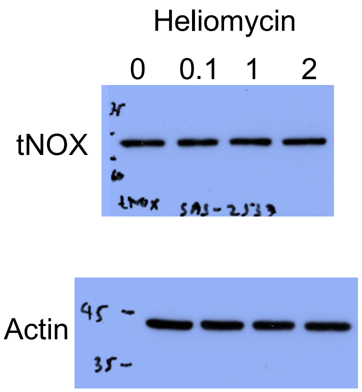


HSC-3

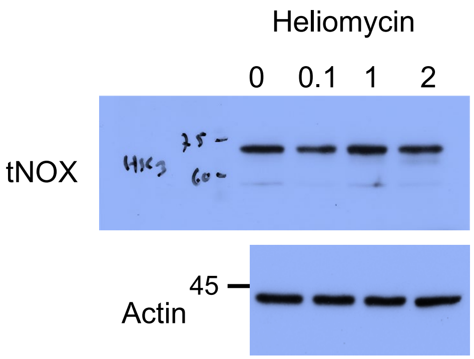

Figure 7 (c): SAS

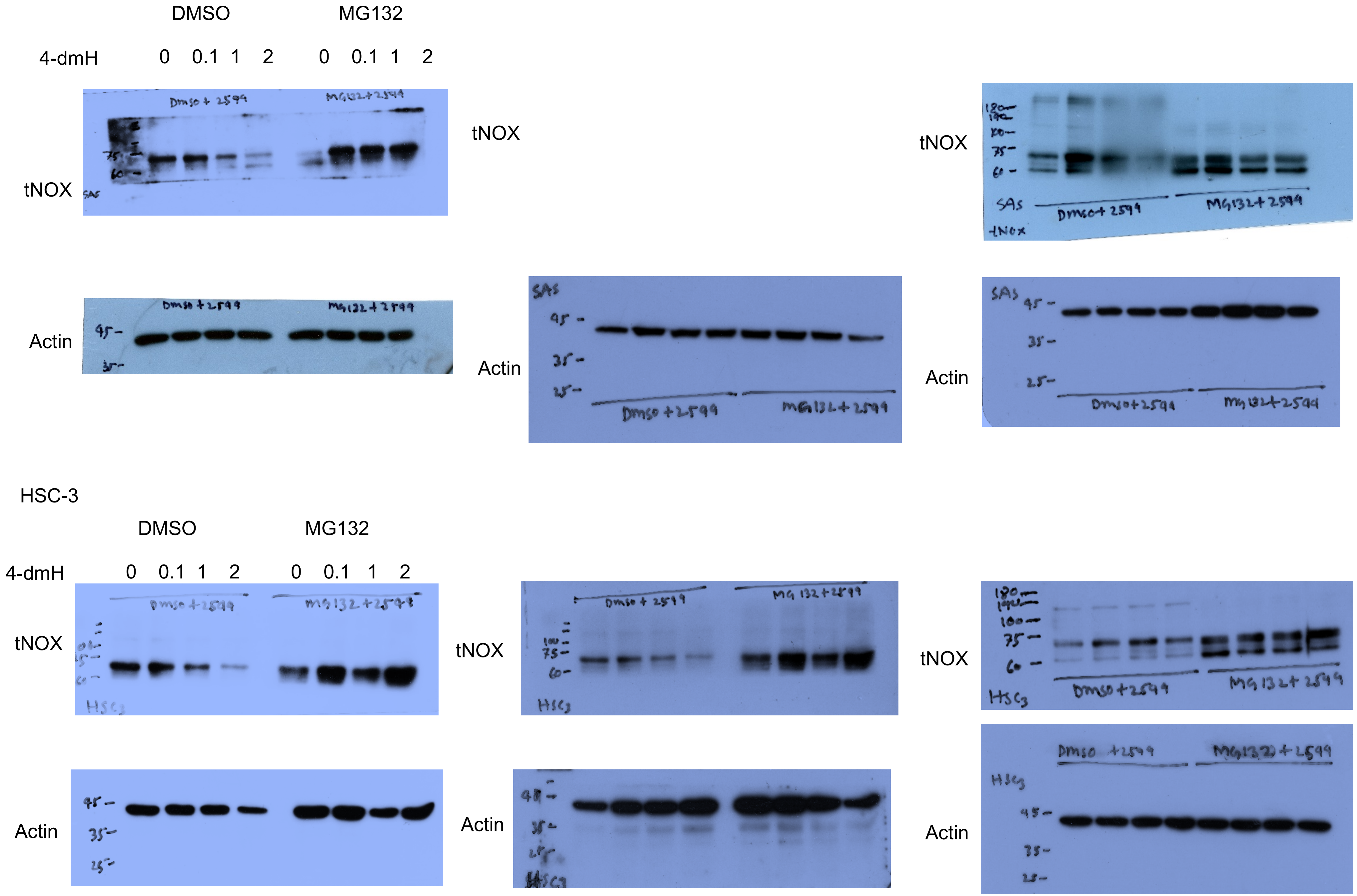

Figure 7 (d): SAS

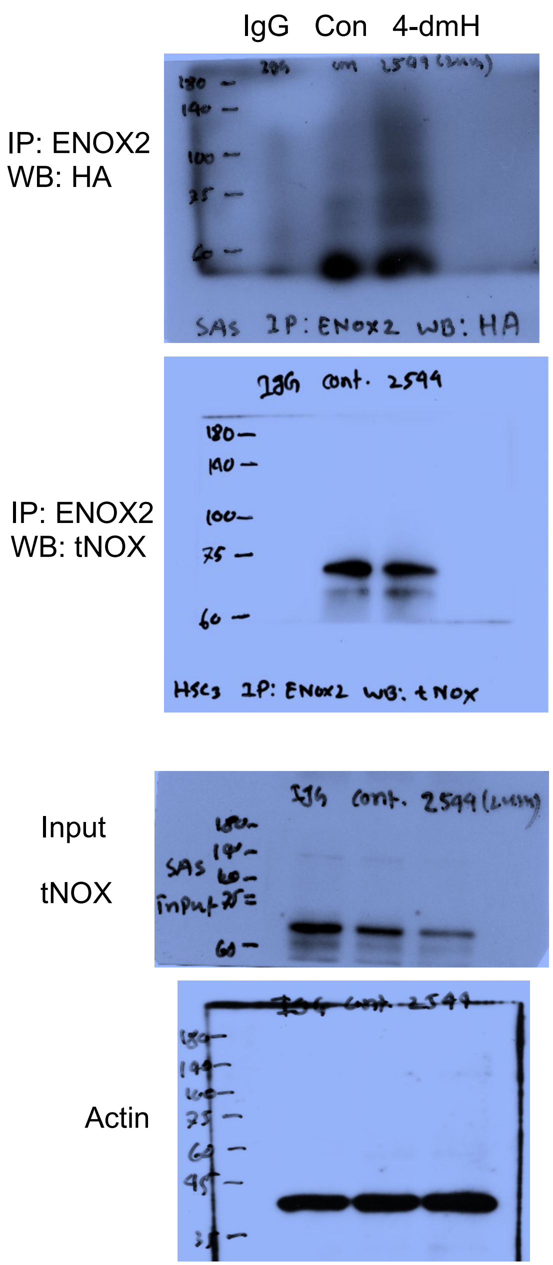


Figure 7 (d): HSC-3

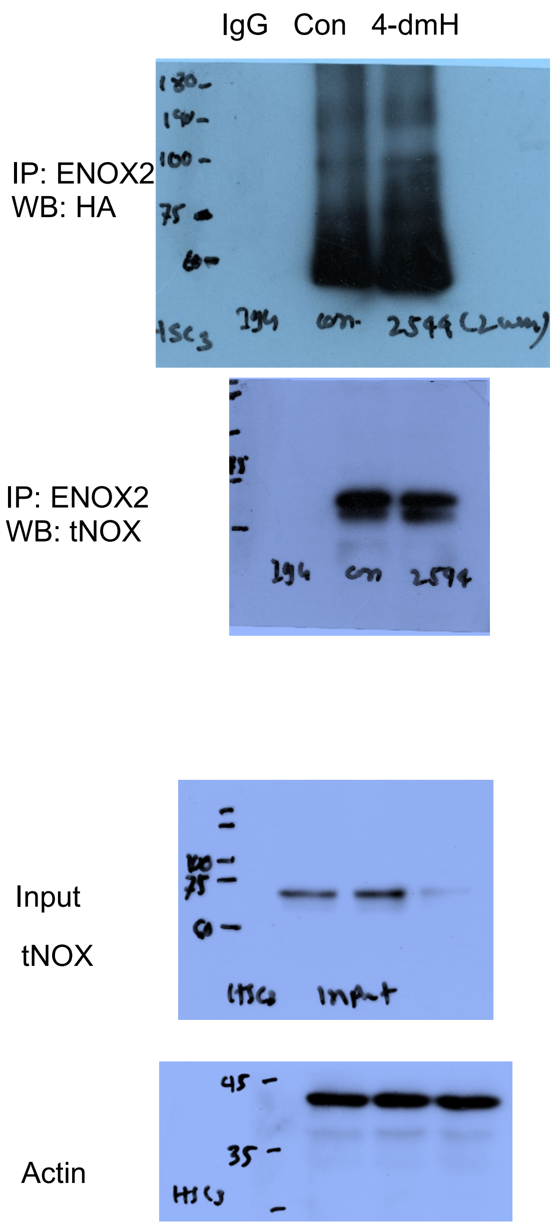


Figure 7 (e)

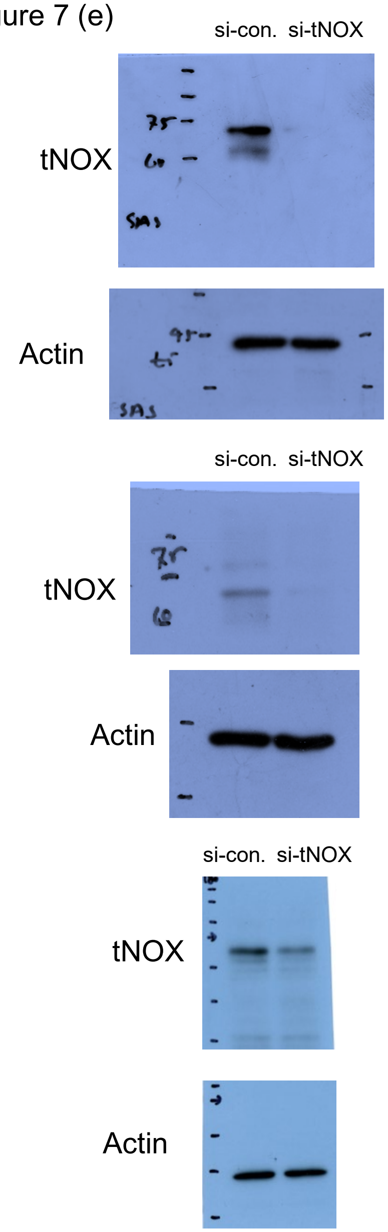

Figure 8 (a): SAS

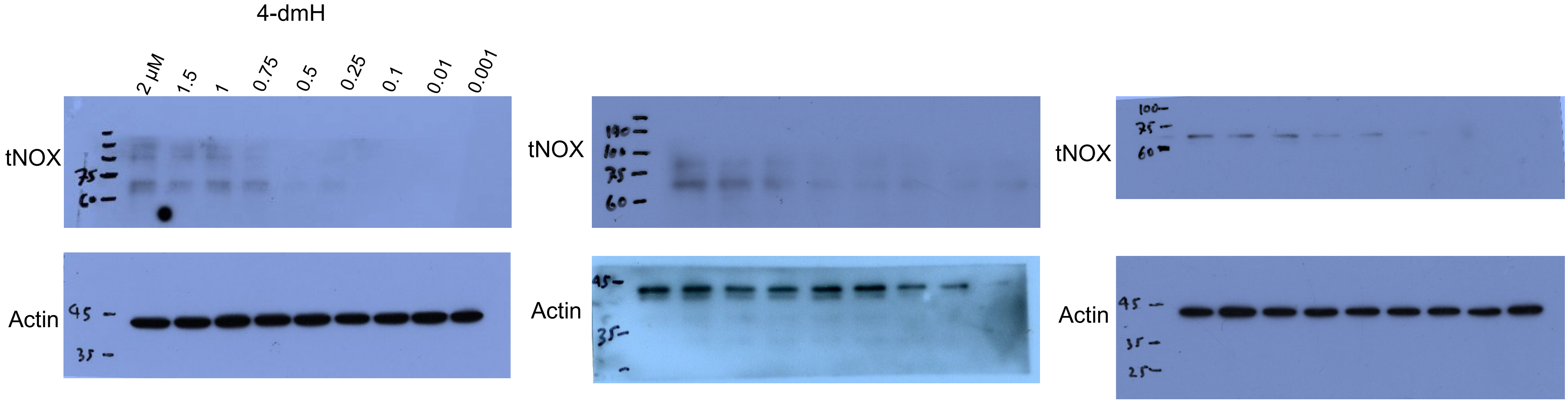


HSC-3

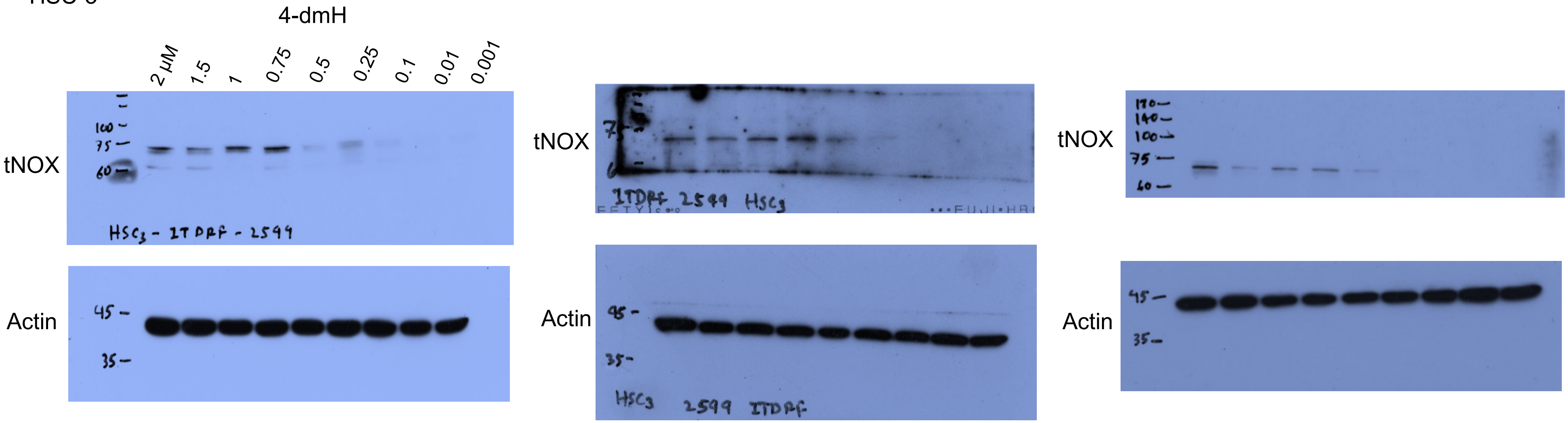

Figure 8 (b): HSC-3

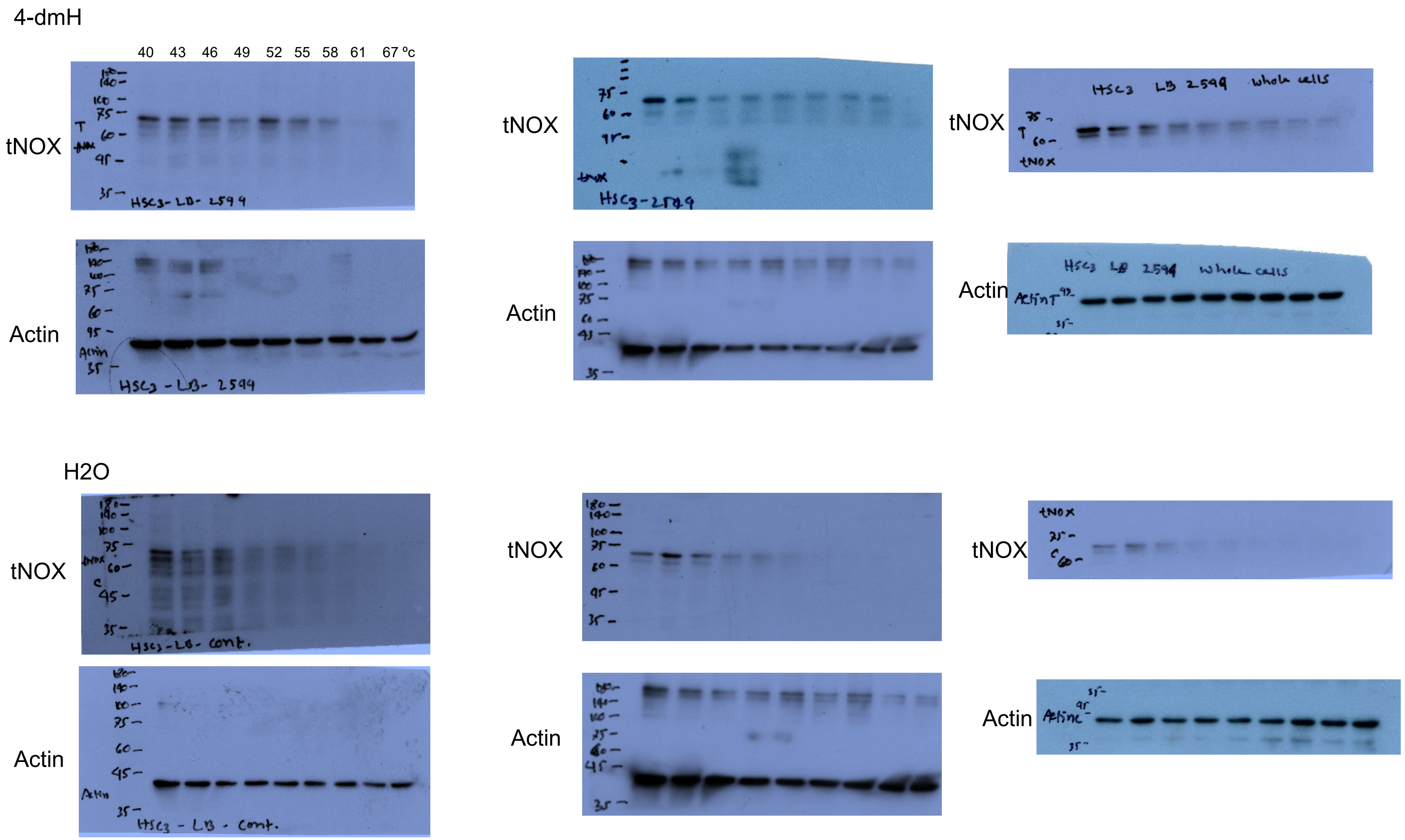

Figure 8 (b): SAS

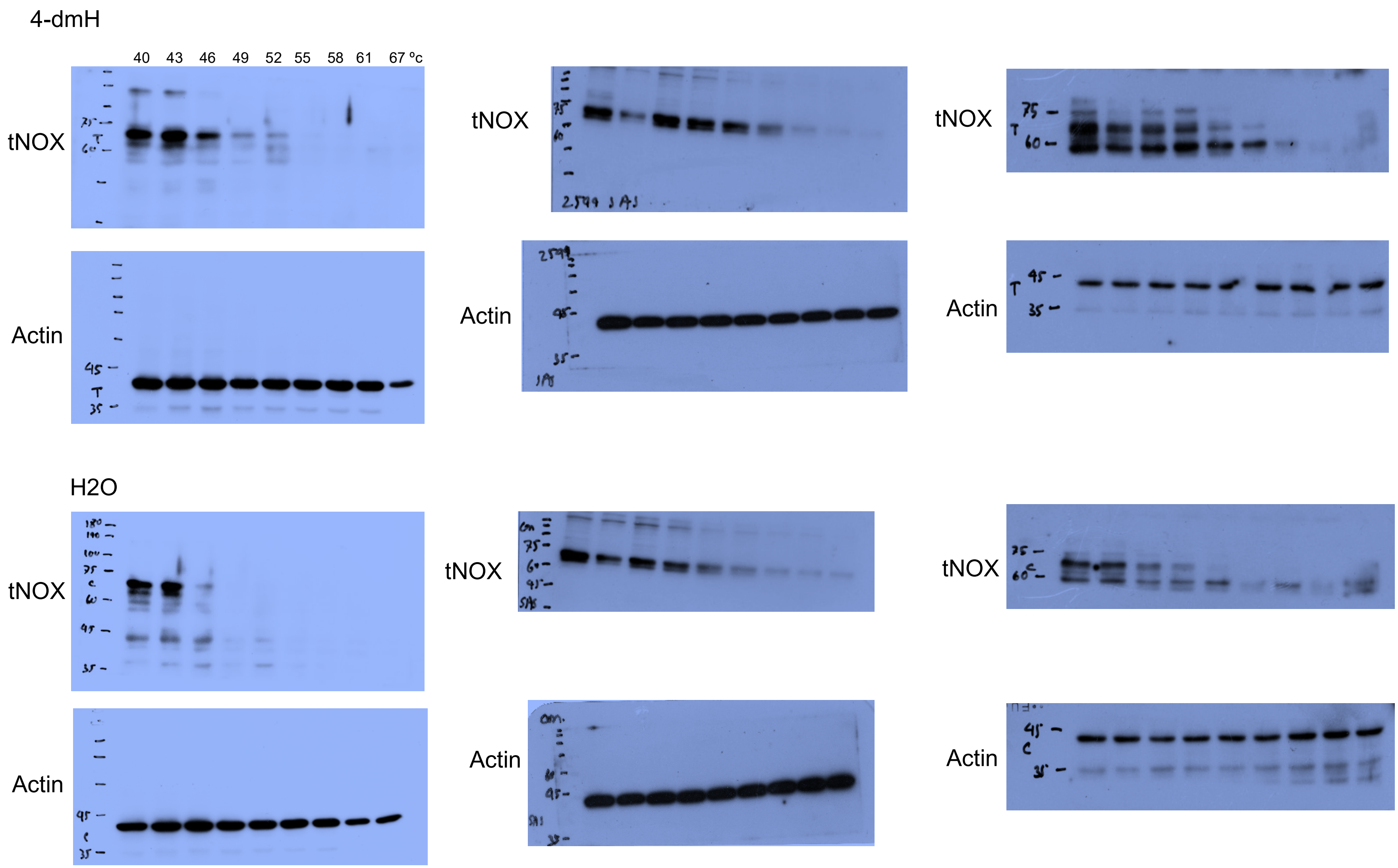

Figure 8 (c): SAS

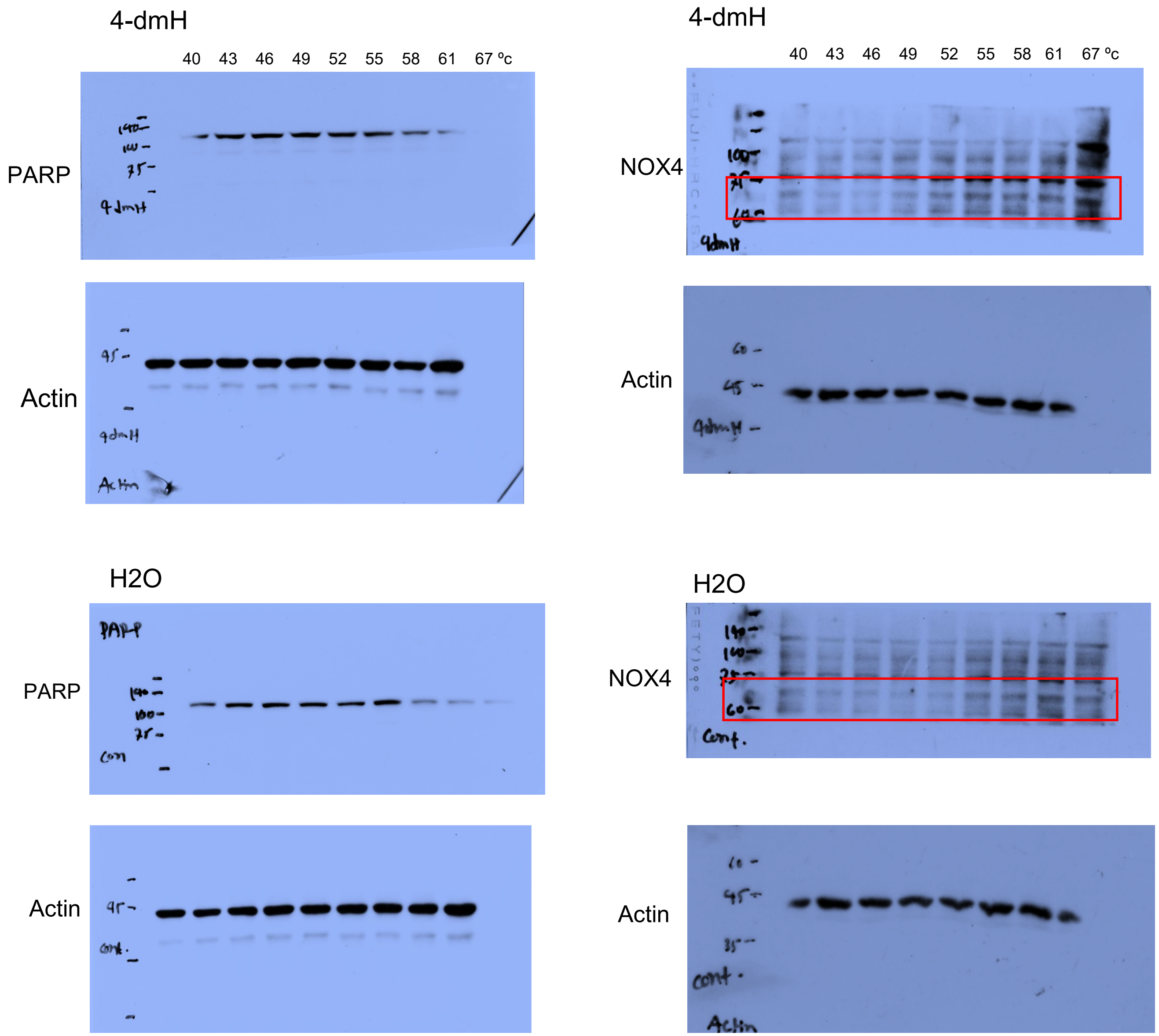

Figure 8 (d): HSC-3

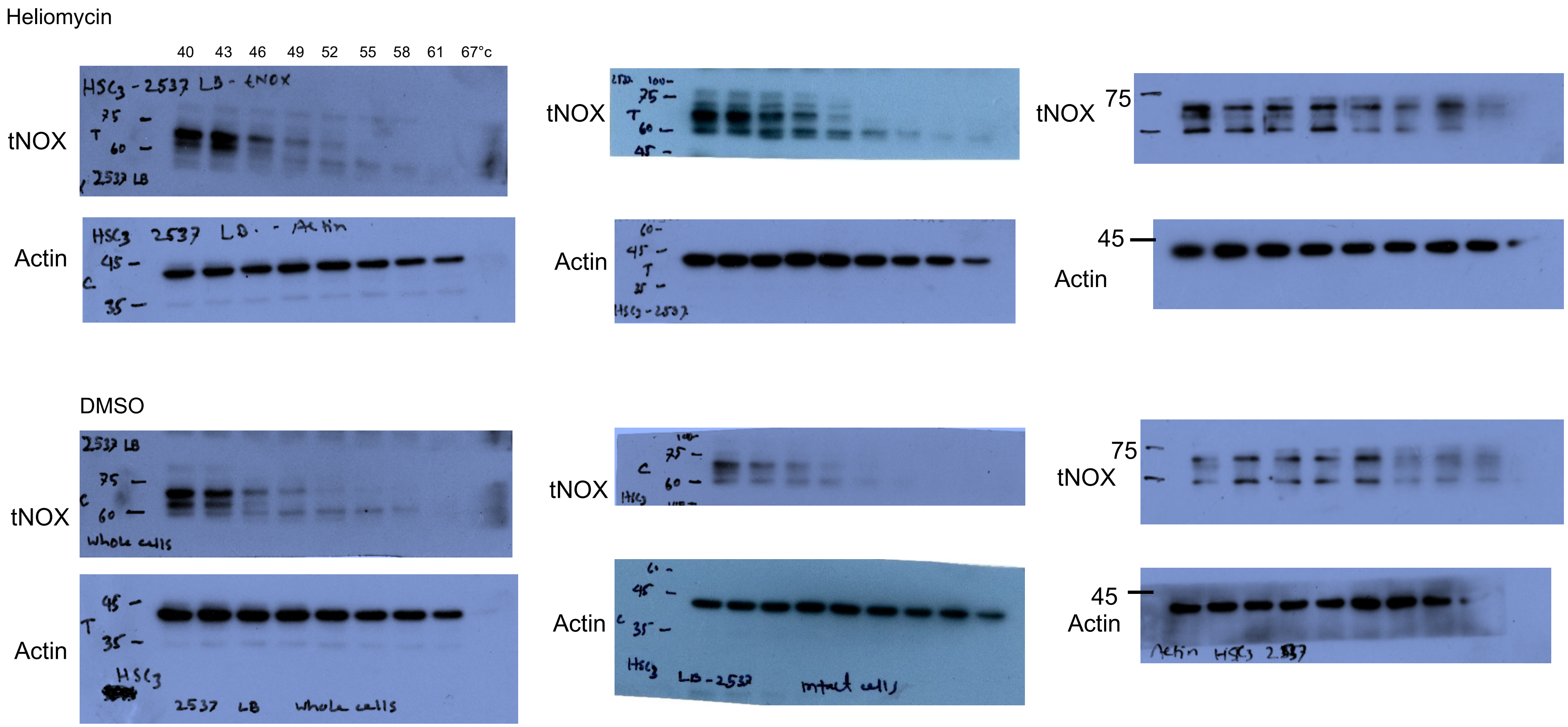

Figure 8 (d): SAS

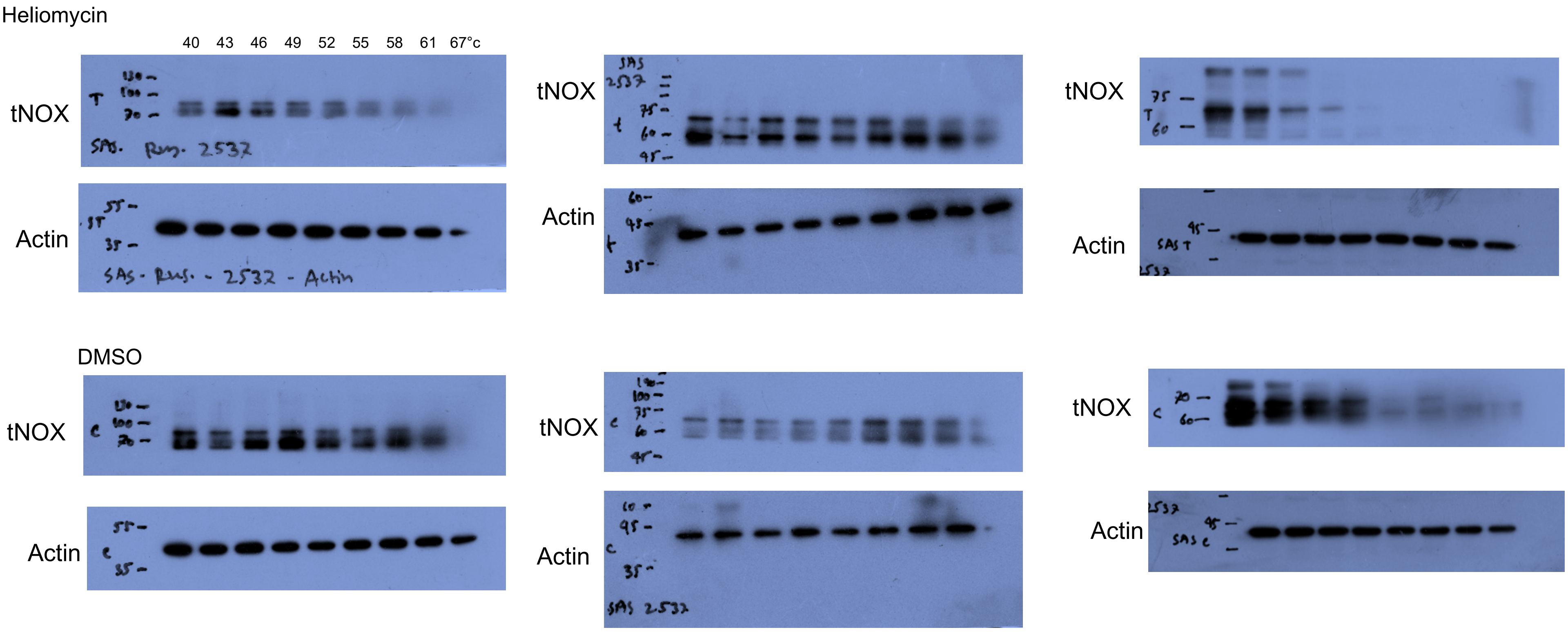

Figure 10 (c): SAS cells inoculated animal tissue samples

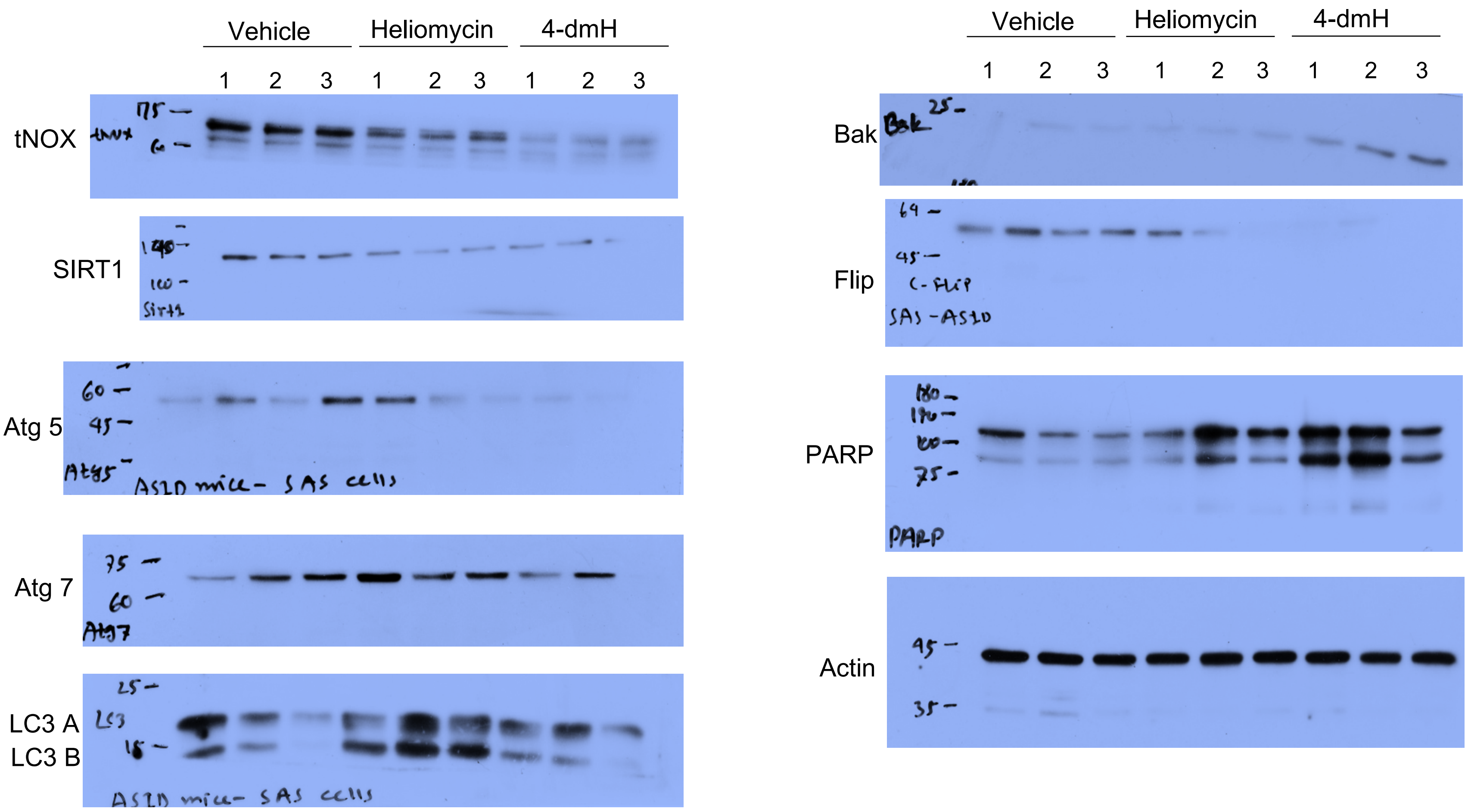

Figure 10 (d)

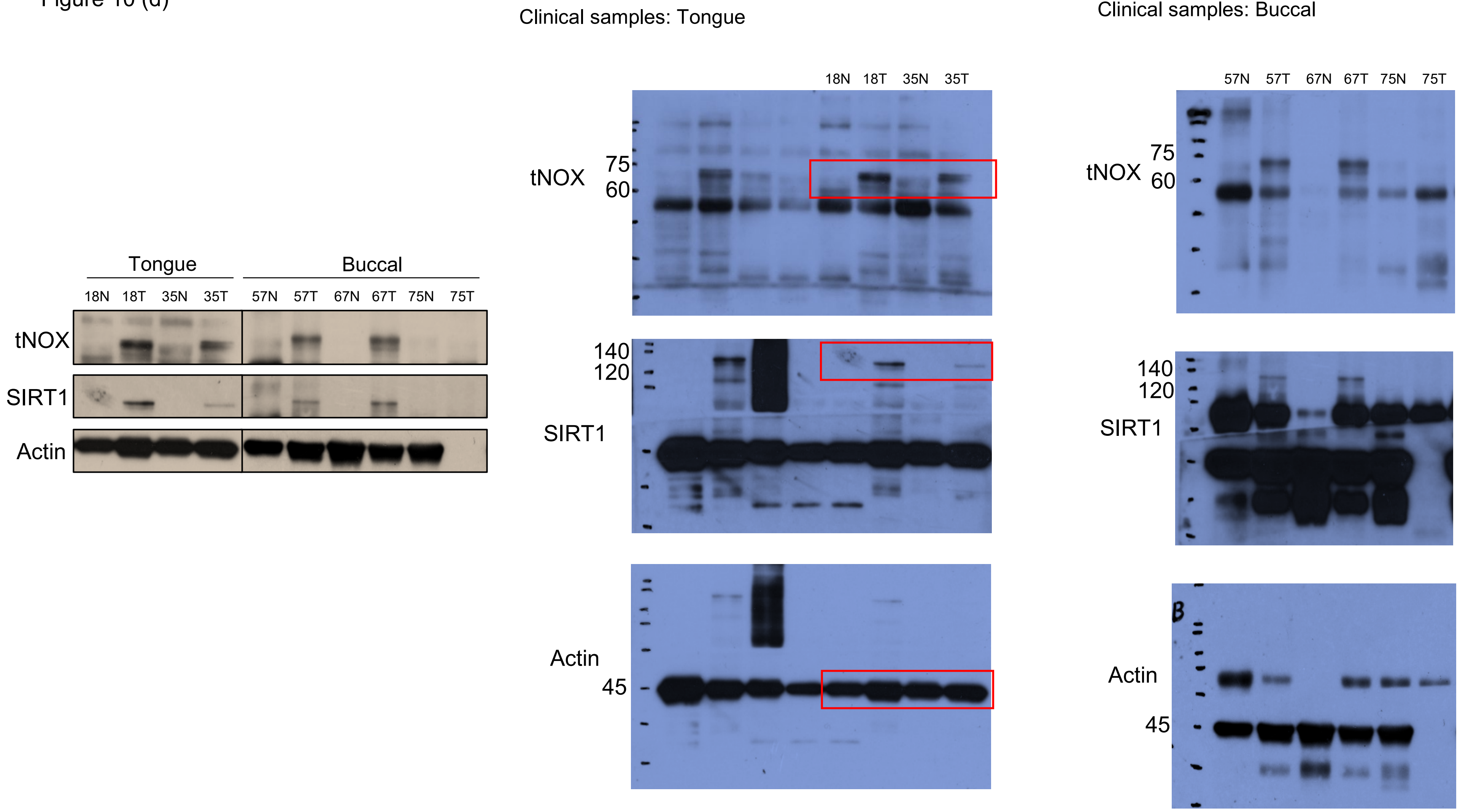
Tongue
Buccal
18N 18T 35N 35T 57N 57T 67N 67T 75N 75T
tNOX
SIRT1
Actin
Clinical samples: Tongue
18N 18T 35N 35T
tNOX
75
60
140
120
SIRT1
Actin
45
Clinical samples: Buccal
57N 57T 67N 67T 75N 75T
tNOX
75
60
140
120
SIRT1
Actin
45
